# Dopaminergic manipulations affect the modulation and meta-modulation of movement speed: evidence from two pharmacological interventions

**DOI:** 10.1101/2023.07.17.549313

**Authors:** Lydia J. Hickman, Sophie L. Sowden, Dagmar S. Fraser, Bianca A. Schuster, Alicia J. Rybicki, Joseph M. Galea, Jennifer L. Cook

## Abstract

A body of research implicates dopamine in the average speed of simple movements. However, naturalistic movements span a range of different shaped trajectories and rarely proceed at a single constant speed; instead, speed is reduced when drawing *“*corners*”* compared to *“*straights*”* (i.e., speed-modulation), and the extent of this slowing down is dependent upon the global shape of the movement trajectory (i.e., speed-meta-modulation) – for example whether the shape is an ellipse or a rounded square. By employing two pharmacological intervention studies – individuals with Parkinson’s both ON and OFF dopaminergic medication (N = 32) and members of the general population on a D2 receptor blocker (haloperidol) versus placebo (N = 43) – we implicate dopamine in speed, speed-modulation and speed-meta-modulation. Our findings move beyond vigour models implicating dopamine in average movement speed, and towards a conceptualisation that involves the modulation of speed as a function of contextual information.

## Introduction

Dopamine is robustly associated with the speed, or ‘vigour’, of movements^1-9^ but naturalistic fluid movements – such as the movements recorded in human handwriting, or when a rat navigates a maze – do not simply proceed at a single, constant, speed. Rather, humans and non-human animals continuously modulate speed according to the curvature of their movements: speeding up along straights and slowing down for corners. ^10, 11^ This phenomenon of adapting movement speed to curvature is mathematically described by a set of scale-invariant power laws^11^ and is robust across species^12-14^ and effectors^10, 15-17^. Speed-modulation of this sort is thought to be a fundamental principle of biological motion but the role of dopamine in speed-modulation is unknown.

Recent advances have shown that speed can also be said to be ‘meta-modulated’. ^11^ That is, the *extent to which* one slows down for corners and speeds up for straights is dependent on the global shape of one’s movement trajectory: If you were drawing a rounded square you would barely modulate your speed (i.e., your speed-modulation value – or the gradient of the slope between your movement speed and current curvature – would be low), whereas speed is dramatically modulated when drawing shapes such as ellipses with fewer, and tighter, ‘corners’. ^11, 18-21^ This means that you would adapt your speed differently to a specific curvature value based on the global shape you are drawing. This observation – that the number of corners in a shape influences the degree to which one slows down for said corners and speeds up for straights – has been mathematically formalised as a spectrum of power laws^11^. If a ‘corner’ is defined as a curvature oscillation per two π of angular displacement, then a shape with two ‘corners’ (an ellipse) is defined as having an angular frequency of two, a shape with four corners (a rounded square) has an angular frequency of four, and so on. Speed-modulation (the gradient of the slope between speed and curvature) is modulated as a function of angular frequency – the higher the angular frequency the lower the speed-modulation.

Although fluid, naturalistic movements nearly always require both speed-modulation and speed-meta-modulation, little is known of the role of dopamine in either of these processes. A number of existing theories link dopamine and movement speed^22-25^, however they do not paint a clear picture of the relationship between dopamine, speed-modulation and speed-meta-modulation. The opportunity costs model^22^, for example, proposes that (tonic) dopamine signals average reward availability, with increased dopamine signalling higher reward availability and thus enhancing the vigour of movements by increasing the opportunity cost of sloth. This model predicts that movements will be less vigorous under low dopamine conditions (e.g., OFF Parkinson’s medication, or under haloperidol – a dopamine antagonist). In line with this, many studies have shown less vigorous movement – long reaction times on button press paradigms^26^ and slow simple reaching arm movements^5, 7, 23^ – under low dopamine conditions. It is, however, unclear how this translates to more complex, naturalistic movements. Consider the example of drawing an ellipse: the opportunity cost of sloth is uniform and unrelated to curvature or global trajectory - it is no more costly to move slowly at corners versus straights. Similarly, it is no more costly to move slowly for low angular frequency (e.g., an ellipse) compared to high angular frequency (e.g., a rounded square) shapes. Therefore, it could be argued that the opportunity costs model^22^ would predict that low dopamine conditions should result in a uniform reduction in the average speed of movement with no change to speed-modulation or speed-meta-modulation (i.e., no modulation as a function of curvature or global trajectory). On the other hand, one could argue that curvature and vigour are related such that straights require more vigorous movements than corners (because straight movements are executed at a higher speed^11, 18-21^), and shapes with more, and less tight, corners (high angular frequency shapes) require more vigorous movement in general (because they tend to be drawn faster^18^). Thus, low dopamine conditions, which reduce vigour, might disproportionately affect straights and high angular frequency shapes because the requisite movements demand more vigour. Under this interpretation, in addition to the effects on speed, the opportunity costs model predicts an effect of dopaminergic manipulation on speed-modulation and speed-meta-modulation.

Existing models, therefore, do not provide clear, unequivocal predictions of the effects of dopaminergic drugs, or of naturally occurring disruptions of the dopamine system (as in Parkinson’s), on natural movements that include speed-modulation and speed-meta-modulation. Here, we investigate the role of dopamine in speed, speed-modulation and speed-meta-modulation. We do so by employing a shapes tracing task in two pharmacological intervention studies in which we study people with Parkinson’s both ON and OFF dopaminergic medication and members of the general population on a D2 receptor blocker (haloperidol) versus placebo.

## Results

Participants comprised individuals with Parkinson’s both ON and OFF dopaminergic medication (Study 1; N=32) and members of the general population on a D2 receptor blocker (haloperidol) versus placebo (Study 2; N=43). In both studies, participants completed a tracing task in which they used a stylus and touch screen device to trace a range of shapes (Fig. 1) for 10 full cycles (where a cycle is a complete start point to start point trace of the shape) per trial. The shapes were strategically chosen on the basis that they span a wide spectrum of angular frequencies and thus enable us to index differences in speed-modulation across the angular frequency spectrum. For each trial of the experiment, we recorded x and y coordinates and calculated indices of speed, speed-modulation (the gradient of the relationship between speed and curvature) and speed-meta-modulation (the gradient of the relationship between speed-modulation and angular frequency). Analyses in the main text were conducted on subsets of the data in which participants had valid trials for at least the shapes with the highest and lowest angular frequency values (Study 1: 4/5 and 4; Study 2: 4/3 and 4). Analyses conducted on full datasets are reported in Supplementary Materials A-D. In the subsequent sections we report dopaminergic effects on speed, speed-modulation and speed-meta-modulation, combining insight from both studies.

**Figure 1.**
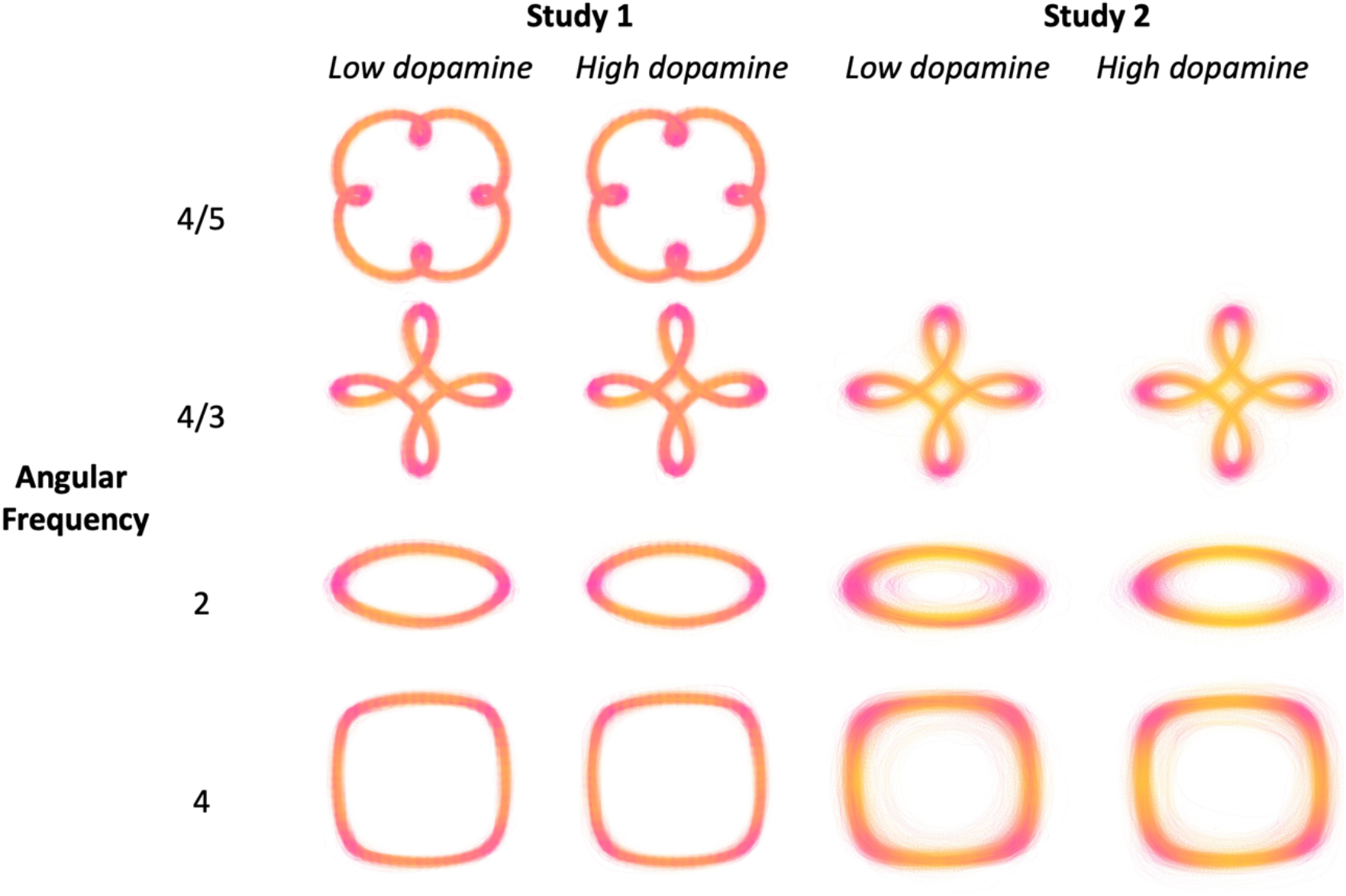
Raw Movement Trajectory Data. Raw trajectory data for all participants under low (Study 1 = Parkinson’s OFF drug; Study 2 = haloperidol) and high (Study 1 = Parkinson’s ON drug; Study 2 = placebo) dopamine conditions in Study 1 (*left*) and Study 2 (*right*), for all trials. The colour scheme represents speed from dark pink (low speed) to yellow (high speed).

### Removal of Parkinson’s medication and administration of haloperidol reduces movement speed

Parkinson’s participants OFF (versus ON) their medication (Study 1; N=32) moved more slowly. That is, a linear mixed model (LMM) including drug state, shape and dosage as fixed effects revealed a main effect of drug state on movement speed (*F*(1,1885)=75.22, *p*<.001; Fig. 2) with lower speed observed OFF medication (beta estimate=-0.081, 95% confidence interval (CI) [-0.099,-0.062]). There was an additional main effect of shape on movement speed (as is typical for shapes across the angular frequency spectrum^18^; *F*(3,1885)=139.41, *p*<.001; Supp Mats A). However, there was no interaction between drug state and shape (indicating that the effect of the drug did not vary as a function of shape and thus it cannot be the case that shapes traced with higher movement speeds were disproportionally affected by the drug), nor an interaction between drug state and dosage (all *p*>.05; Supp Mats A).

**Figure 2.**
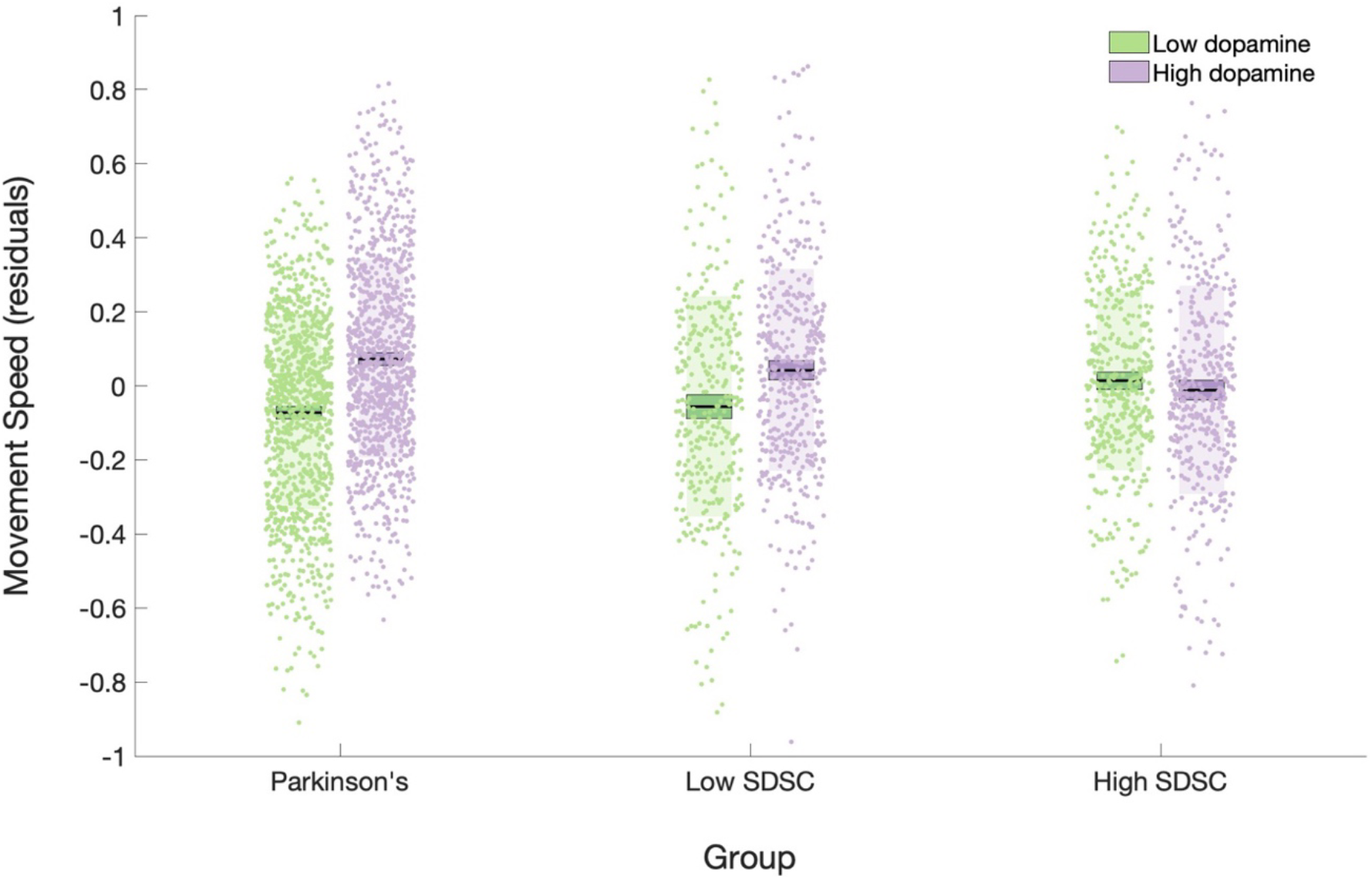
A Graph Depicting Drug Effects on Movement Speed. Movement speed residuals plotted for low (green) and high (purple) dopamine conditions across 3 groups: individuals with Parkinson’s OFF and ON their dopaminergic medication (Study 1); members of the general population with low striatal dopamine synthesis capacity (as determined by a median split) under haloperidol or placebo (Study 2); members of the general population with high striatal dopamine synthesis capacity under haloperidol or placebo (Study 2). Residuals were calculated after controlling for dosage (Study 1) or striatal dopamine synthesis capacity (Study 2), and trial, participant number and day (Studies 1 and 2). Main effects of drug state revealed slower movement speed in low dopamine conditions, with an interaction between drug and striatal dopamine synthesis capacity in Study 2 indicating opposing drug effects in the two groups. Bars = mean, box = SEM, shaded region = standard deviation, trial-level values plotted, SDSC = striatal dopamine synthesis capacity.

Similarly, we found that administration of the dopamine D2 receptor blocker haloperidol to members of the general population (Study 2; N=43) reduced movement speed. An LMM including drug state, shape and estimated baseline striatal synthesis capacity as fixed effects revealed a significant main effect of drug state (*F*(1,1600)=45.43, *p*<.001) with lower movement speed values in the haloperidol condition compared to the placebo condition (beta estimate: -0.523, 95%CI[-0.676,-0.371]). Again, a main effect of shape (*F*(2,1600)=175.24, *p*<.001; Supp Mats B) was identified, and no interaction between drug state and shape (p>.05; Supp Mats B), again highlighting that the drug effect was not disproportionally higher for shapes traced at higher movement speeds. Results from both studies indicate slowed movement under low dopamine conditions.

A body of literature reports that effects of dopaminergic drugs are modulated by striatal dopamine synthesis capacity^27-29^ and that working memory capacity can serve as a proxy for this, with low working memory capacity indicating low dopamine synthesis capacity^30, 31^. This literature argues for an inverted-U function for the relationship between baseline dopamine and the effects of dopaminergic drugs on performance^32^. To enable us to explore any baseline dependent effects of the drug we asked participants in Study 2 to complete a working memory task^33^ whilst under placebo. We predicted that, if there is an inverted-U-shaped relationship between dopamine and speed we would see that the effect of haloperidol is moderated by estimated striatal dopamine synthesis capacity such that individuals with low striatal dopamine synthesis capacity move slower under haloperidol relative to placebo because their (already sub-optimally low) dopamine levels are further reduced by haloperidol. In contrast, individuals with high striatal dopamine synthesis capacity should show speeding effects of the drug because their sub-optimally high levels of dopamine are reduced by haloperidol, thus bringing them closer to the optimal level of dopamine for speedy movements. Including our proxy for striatal dopamine synthesis capacity (i.e., working memory score) as a covariate in our mixed model revealed that the main effect of drug state was indeed moderated by estimated striatal dopamine synthesis capacity (*F*(1,1600)=42.04, *p*<.001; Fig. 2).

The interaction between drug state and estimated striatal dopamine synthesis capacity was unpacked by calculating the drug effect for each participant (mean speed under placebo minus mean speed under haloperidol) and plotting this against estimated baseline striatal dopamine synthesis capacity. Results are plotted in Fig. 5 (*top-left*). Positive values on the y-axis indicate that participants’ movements were slowed by haloperidol administration, and negative values indicate that participants moved faster under haloperidol than placebo. Our data revealed a negative linear relationship between drug effect and estimated striatal dopamine synthesis capacity which cut the x-axis, indicating opposing drug effects in low versus high striatal dopamine synthesis capacity groups. That is, a haloperidol-induced reduction in dopamine caused participants with low striatal dopamine synthesis capacity to move slower, and those with high striatal dopamine synthesis capacity to move faster. These results suggest that the drug effect on speed is dependent on estimated striatal dopamine synthesis capacity.

### Removal of Parkinson’s medication and administration of haloperidol reduces speed-modulation

To investigate effects on speed-modulation we carried out LMMs with speed-modulation values – the gradient of the regression line between instantaneous movement speed and the curvature being drawn – as the dependent variable. Here a main effect of drug state would comprise evidence that manipulating dopamine impacts speed-modulation. An interaction between drug state and shape would be evidence of a drug effect on *speed-meta-modulation;* as such, in this section we focus on main effects and discuss interactions between drug state and shape in the next section.

In addition to effects on speed, we observed that removal of Parkinson’s medication (Study 1) affected speed-modulation. Our LMM with drug state, shape and dosage as fixed effects revealed a main effect of drug state on speed-modulation (*F*(1,1906)=8.12, *p*=.004; Fig.3), whereby lower speed-modulation values were observed OFF Parkinson’s medication (beta estimate=-0.017, 95%CI[-0.029,-0.005]; Supp Mats C for main effect of shape). No interaction between drug state and dosage was found (*p*>.05; Supp Mats C).

A comparable analysis with members of the general population (Study 2) revealed a parallel drug effect on speed-modulation. That is, a LMM with drug state, shape and estimated baseline striatal dopamine synthesis capacity as fixed effects and speed-modulation as the dependent variable, revealed a significant main effect of drug (*F*(1,1642)=35.68, *p*<.001). Consistent with Study 1, speed-modulation values were lower under haloperidol compared to placebo (beta estimate=-0.233, 95%CI[-0.310,-0.157]). The interaction between drug state and estimated striatal dopamine synthesis capacity was also significant (*F*(1,1642)=42.85, *p*<.001; Fig. 3; Supp Mats D for main effect of shape), again illustrating the strong baseline-dependency of the effect. Fig. 5 (*top-right*) illustrates a significant negative linear relationship between the drug effect and estimated striatal dopamine synthesis capacity (*F*(1,22)=6.96, *p*=.015; beta estimate=-0.006, 95%CI[-0.012,-0.001]) which again cuts the x-axis, indicating opposing drug effects in low versus high striatal dopamine synthesis capacity groups. That is, a reduction in dopamine caused participants with low striatal dopamine synthesis capacity to move with reduced speed-modulation, and those with high striatal dopamine synthesis capacity to move with increased speed-modulation. Thus, as is the case for speed, the drug effect on speed-modulation is dependent on estimated striatal dopamine synthesis capacity.

**Figure 3.**
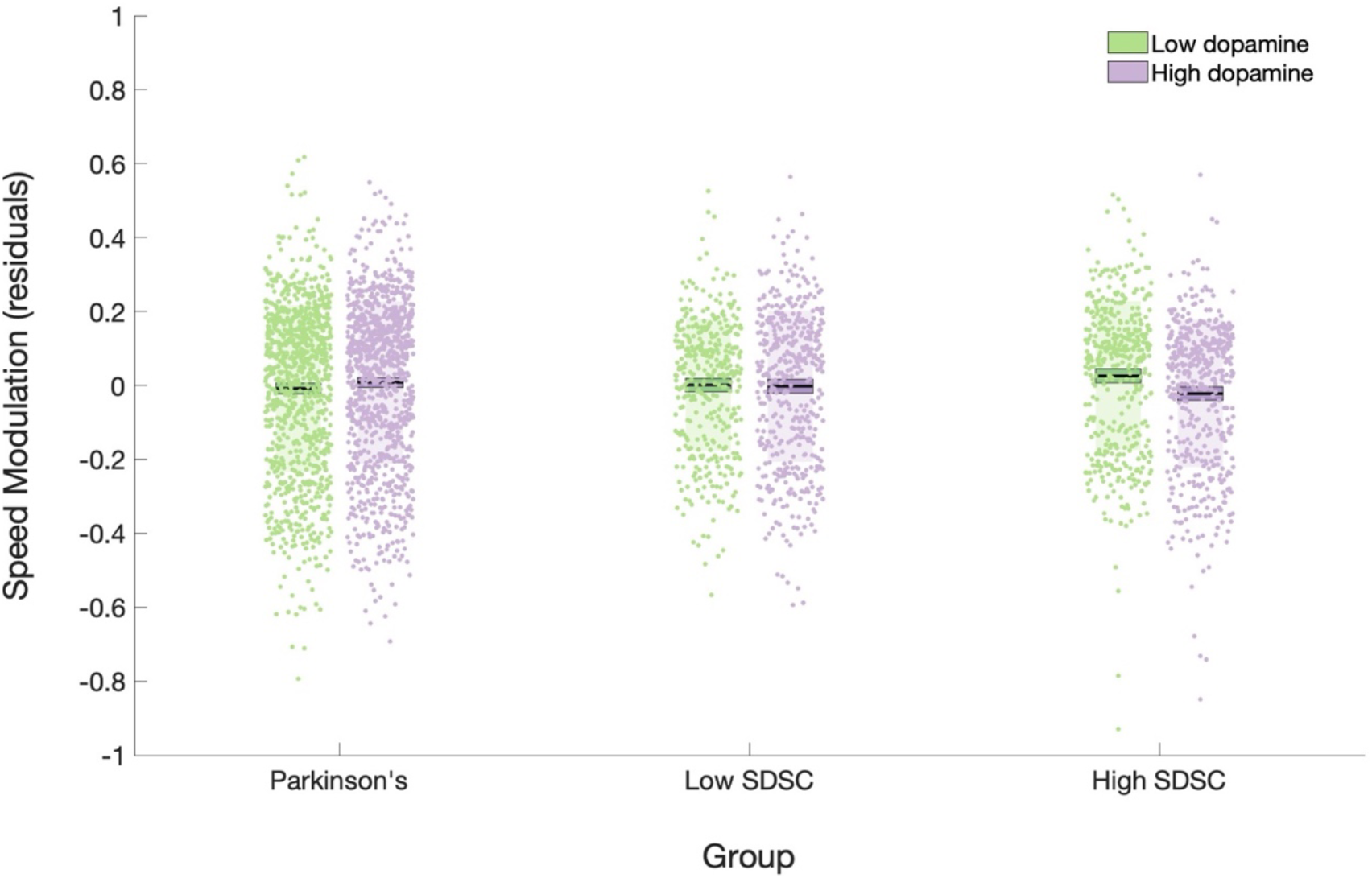
A Graph Depicting Drug Effects on Speed-Modulation. Speed-modulation residuals plotted for low (green) and high (purple) dopamine conditions across 3 groups: individuals with Parkinson’s OFF and ON their dopaminergic medication (Study 1); members of the general population with low striatal dopamine synthesis capacity (as determined by a median split) under haloperidol or placebo (Study 2); members of the general population with high striatal dopamine synthesis capacity under haloperidol or placebo (Study 2). Residuals were calculated after controlling for dosage (Study 1) or striatal dopamine synthesis capacity (Study 2), and trial, participant number and day (Studies 1 and 2). Main effects of drug state revealed reduced speed-modulation values in low dopamine conditions, with an interaction between drug and striatal dopamine synthesis capacity in Study 2 indicating opposing drug effects in the two groups. Bars = mean, box = SEM, shaded region = standard deviation, trial-level values plotted, SDSC = striatal dopamine synthesis capacity.

**Figure 4.**
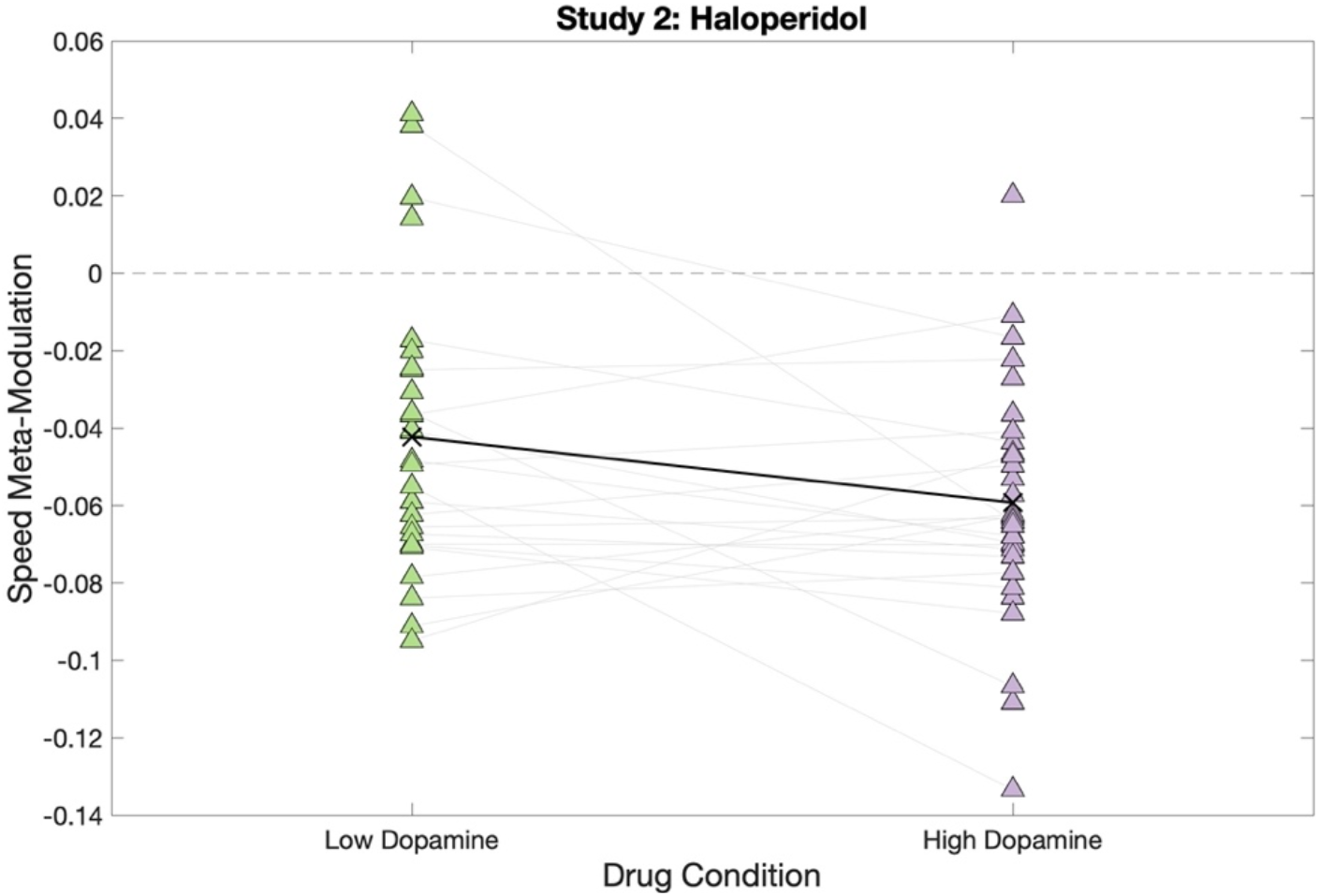
A Graph Depicting Drug Effects on Speed-meta-modulation. Speed-meta-modulation values for each participant in low (haloperidol; green) versus high (placebo; purple) dopamine conditions for Study 2. A main effect of drug state was observed, revealing lower speed-meta-modulation (closer to zero) in low dopamine conditions. Light grey lines represent participant data; black crosses indicate mean values for each condition.

**Figure 5.**
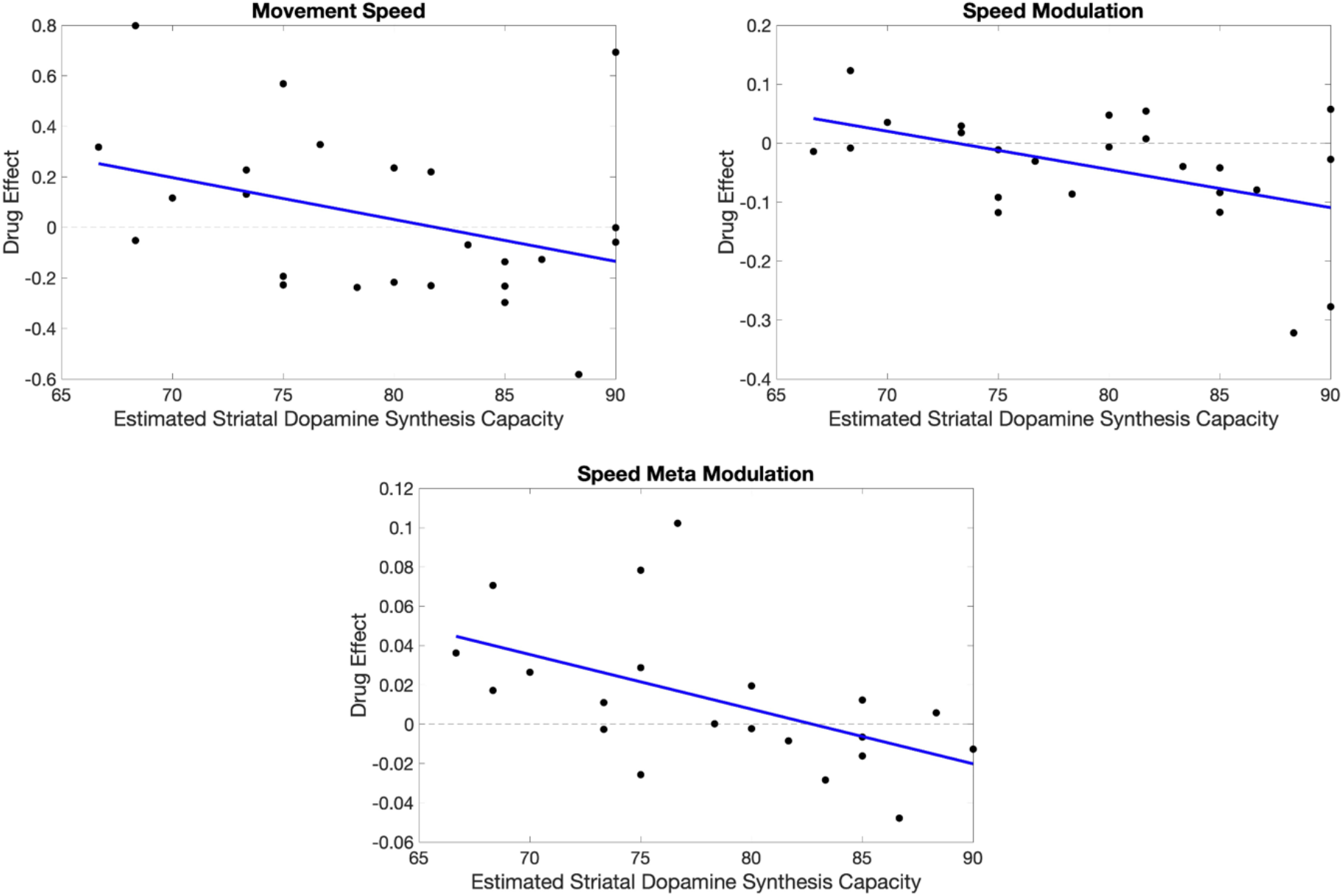
Graphs Depicting Striatal Dopamine Synthesis Capacity Dependency of Drug Effects. Each graph presents the drug effect (calculated as the difference in dependent variable between low and high dopamine conditions) per participant in Study 2 plotted against the estimated striatal dopamine synthesis capacity for that participant. Graphs refer to movement speed (top-*left*), speed-modulation (*top-right*) and speed-meta-modulation (*bottom*) respectively. In each case, low dopamine conditions reduce performance on the dependent variable in those with low striatal dopamine synthesis capacity, and increase it with high striatal dopamine synthesis capacity, as indicated by a linear trend which cuts the x-axis. Blue line = line of best fit.

**Table 1.** Drug Effects for Speed, Speed-Modulation and Speed-Meta-Modulation.

| | $\beta$ estimate | SE | Lower CI | Upper CI | F statistic | DoF | p value |
| --- | --- | --- | --- | --- | --- | --- | --- |
| **Speed** |  |  |  |  |  |  |  |
| Study 1: Parkinson's Medication | -0.081 | 0.01 | -0.099 | -0.062 | 75.22 | 1, 1885 | <.001*** |
| Study 2: Haloperidol | -0.523 | 0.08 | -0.676 | -0.371 | 45.43 | 1, 1600 | <.001*** |
| **Speed-Modulation** |  |  |  |  |  |  |  |
| Study 1: Parkinson's Medication | -0.017 | 0.01 | -0.029 | -0.005 | 8.12 | 1, 1906 | .004** |
| Study 2: Haloperidol | -0.233 | 0.04 | -0.310 | -0.157 | 35.68 | 1, 1642 | <.001*** |
| **Speed-Meta-Modulation** |  |  |  |  |  |  |  |
| Study 1: Parkinson's Medication | 0.002 | 0.00 | -0.004 | 0.008 | 0.49 | 1, 57 | .485 |
| Study 2: Haloperidol | 0.107 | 0.03 | 0.046 | 0.167 | 12.40 | 1, 53 | .001** |

*Note.* \* $p < .05$ , \*\* $p < .01$ , \*\*\* $p < .001$ . SE = standard error, DoF = degrees of freedom, CI = confidence interval.

Our Introduction highlighted that a possible prediction from the opportunity costs model is that a reduction in vigour following dopaminergic manipulation will disproportionately affect high speed movements that require more vigour. This would predict a greater effect of the drug on parts of the trajectories that are executed at higher speeds (i.e., straights relatives to corners), and may account for reductions in speed-modulation values. To interrogate the data for evidence in support of this, we analysed the average of the top and bottom 10% of speed values obtained during each trial using the same LMM designs as set out previously for movement speed. For Study 1 there was no significant effect of drug state on either maximum or minimum speed values (maximum values: *F*(1,1892)=1.03, *p*=.309; minimum values: *F*(1,1955)=0.00, *p*=.984). For Study 2, whilst there was no drug effect on maximum speed values (*F*(1,1612)=3.21, *p*=.073), minimum speed values were significantly higher under haloperidol (*F*(1,1637)=12.63, *p*<.001; beta estimate=0.074, 95%CI[0.033,0.114]), resulting in a reduced range of speed values. Thus, neither study provided evidence that dopaminergic manipulation disproportionately affects higher speed movements.

### Administration of haloperidol reduces speed-meta-modulation, but Parkinson’s medication does not

To test whether the removal of Parkinson’s medication (Study 1) would reduce the meta-modulation of movement speed, we calculated a speed-meta-modulation index by regressing angular frequency against speed-modulation values and calculating the gradient of the regression line. Here, a larger gradient magnitude would indicate a greater difference in speed-modulation across the angular frequency spectrum. This variable was submitted to a LMM with drug state and dosage as fixed effects. There was no significant effect of drug state on speed-meta-modulation, nor drug state by dosage interaction (*p*>.05; Supp Mats E). This is supported by the lack of a drug state by shape interaction in the speed-modulation LMM described above for Study 1 (*p*>.05; Supp Mats E). As such, in Study 1 we did not find evidence supporting a role for dopamine in speed-meta-modulation.

By contrast, in the general population (Study 2) we observed that haloperidol reduced the meta-modulation of speed (*F*(1,53)=12.40, *p*=.001; beta estimate=0.107, 95%CI[0.046,0.167]), whereby speed-meta-modulation gradients were flatter (i.e., had a lower magnitude or were ‘less negative’) under haloperidol. This indicated that, under haloperidol, participants did not utilise a range of speed-modulation values across the angular frequency spectrum as would be seen in appropriate speed-meta-modulation. Instead, the extent to which speed was modulated as a function of curvature was similar across all shapes, thus reducing the gradient of the relationship between angular frequency shape and speed-modulation value. This is supported by the presence of a drug state by shape interaction in the speed-modulation LMM described above for Study 2 (*F*(1,1642)=11.45, *p*<.001). Therefore, in Study 2, we find evidence for reduced speed-meta-modulation in low dopamine conditions.

As was the case for movement speed and speed-modulation, the effect of haloperidol on speed-meta-modulation was baseline dopamine dependent; a significant interaction between drug state and estimated striatal dopamine synthesis capacity (*F*(1,53)=10.95, *p*=.002) was observed. To aid interpretation, we plotted the relationship between the drug effect and striatal dopamine synthesis capacity (Fig. 5, *bottom*). The drug effect (y-axis) was calculated as the speed-meta-modulation value under haloperidol minus the speed-meta-modulation value under placebo. Note that for speed and speed-modulation (Fig. 5, *top-left* and *top-right*) the drug effect was instead calculated as placebo minus haloperidol because more positive speed and speed-modulation values indicate higher speed and greater speed-modulation. Given that more positive speed-meta-modulation values actually indicate *less* speed-meta-modulation (i.e., flatter slopes for the negative relationship between angular frequency and speed-modulation), the drug effect was calculated as haloperidol minus placebo to ensure that, in line with the other two dependent variables, positive values on the y-axis indicate reduced speed-meta-modulation under haloperidol and negative values indicate increased speed-meta-modulation.

Fig. 5 (*bottom*) demonstrates an effect consistent with the other two dependent variables: a significant negative linear relationship between drug effect and estimated striatal dopamine synthesis capacity (*F*(1,19)=8.31, *p*=.010; beta estimate=-0.003, 95%CI[-0.005,-0.001]). These data again cut the x-axis, indicating opposing drug effects whereby, a reduction in dopamine decreased speed-meta-modulation for participants with low striatal dopamine synthesis capacity, and increased speed-meta-modulation for those with high striatal dopamine synthesis capacity. As such, all three dependent variables exhibit drug effects that are dependent on estimated striatal dopamine synthesis capacity.

Our Introduction highlighted that a possible prediction from the opportunity costs model is that a reduction in vigour following dopaminergic manipulation will disproportionately affect shapes at higher angular frequencies because these tend to be drawn at a higher speed and may therefore require more vigour. To interrogate the data for evidence in support of this we further explored the drug state by shape interaction in the speed-modulation LMM for Study 2 by running the same LMM (excluding shape) on subsets of the data for each shape. This revealed that the ‘flattening of the curve’ between angular frequency and speed-modulation (i.e., reduced speed-meta-modulation) under low dopamine conditions was driven by larger drug effects for the lower angular frequency shapes than the higher angular frequency shapes (shape 4/3: *F*(1,555)=49.36, *p*<.001, beta estimate=-0.391, 95%CI[-0.500,-0.282]; shape 2: *F*(1,549)=14.78, *p*<.001, beta estimate=-0.202, 95%CI[-0.306,-0.099]; shape 4: *F*(1,534)=2.95, *p*=.086, beta estimate=-0.127, 95%CI[-0.273,0.018). Thus, there is no evidence that dopaminergic manipulation disproportionately affected high angular frequency shapes.

### Drug effects on speed, speed-modulation and speed-meta-modulation are independent of each other

Given the presence of drug effects on all three dependent variables in Study 2, further exploratory analyses were implemented on this dataset to investigate whether the drug effects are independent of each other. A main effect of drug remained for speed after controlling for speed-modulation (*F*(1,1569)=34.40, *p*<.001) and speed-meta-modulation (*F*(1,1510)=52.23, *p*<.001). Similarly, the effect of the drug on speed-modulation persisted after controlling for speed (*F*(1,1565)=10.36, *p*=.001) and speed-meta-modulation (*F*(1,1554)=17.66, *p*<.001). Finally, the main effect of drug for speed-meta-modulation remained after controlling for speed (*F*(1,53)=12.41, *p*=.001) and speed-modulation (*F*(1,53)=18.18, *p*<.001). These results indicate that there are separable effects of the drug on speed, speed-modulation and speed-meta-modulation.

## Discussion

In line with the vigour literature, our results showed that low dopamine conditions (removal of Parkinson’s medication and administration of haloperidol relative to placebo) reduced movement speed. In addition, dopaminergic drugs independently affected speed-modulation by reducing participants’ ability to modulate movement speed according to curvature. Finally, we also found an independent effect of dopaminergic drugs on speed-meta-modulation: haloperidol reduced participants’ ability to titrate their speed-modulation such that it was appropriately suited to the global trajectory (the angular frequency of the shape). This latter effect was seen in members of the general population but not for individuals with Parkinson’s; we speculate below that this may be due to the incorporation of estimated baseline striatal dopamine synthesis capacity in the general population study. Together these results implicate dopamine in average movement speed, speed-modulation and speed-meta-modulation, and show that dopamine’s role in movement speed is broader than that which is conceptualised in current models linking dopamine and movement.

Existing models, such as the opportunity costs models, do not provide clear unequivocal predictions for the effects of dopaminergic manipulation on naturalistic movement. Nevertheless, our Introduction highlighted two *possible* predictions that can be made from the opportunity costs model. One prediction is that low dopamine conditions should be associated with speed reductions but with no effects on speed-modulation and speed-meta-modulation. Our results clearly diverge from this prediction and thus show that this interpretation of the opportunity costs model does not align with empirical evidence. A second prediction is that dopaminergic manipulation will affect speed, speed-modulation and speed-meta-modulation because reductions in vigour will disproportionately affect high speed movements that require more vigour. More specifically, this would predict a greater effect of the drug on trajectories that are executed at higher speeds (straights vs. corners) and shapes that are executed at higher speeds (high angular frequency shapes). We did not find any evidence to support these predictions. That is, our data did not show that drug state disproportionately affected higher speed movements, or that drug state disproportionately affected speed or speed-modulation values for high, as opposed to low, angular frequency shapes. Our results, cannot, therefore, easily be conceptualised under current formulations of the opportunity costs model.

We next consider whether our results might be consistent with other popular models in the literature. Bayesian theories^34, 35^ propose that (tonic) dopamine signals the precision with which incoming information is stored and represented. Under high dopamine – signalling high precision – an individual is thought to be more confident in their representation of incoming sensory information compared to their prior beliefs, and thus more heavily relies on incoming sensory data; conversely, low dopamine promotes a reliance on prior beliefs. Whilst Bayesian theories have not made *explicit* predictions about our task, one can consider tracing as a task that requires a balance between priors and incoming evidence. One has priors, learned through previous experience, that influence our typical speed of movement^36^ but as we trace around the outline of a shape we encounter changes in curvature that demand deviations from these priors. Thus, if we assume that reduced speed-modulation and meta-modulation can occur due to an overweighting of the prior (typical speed) relative to the incoming (trajectory) information, then our results could be considered consistent with Bayesian theories of dopamine and movement. Bayesian theories do not, however, provide a comprehensive account of *all* of our results because they do not make clear predictions about average movement speed.

A more recent theory - the rational inattention account^37^ - merges opportunity cost and Bayesian models by proposing that dopamine signals average reward availability and that this ‘pays the cognitive costs’ (e.g., attention costs) of increasing precision. This is consistent with evidence that dopamine plays a role in both motivational and cognitive control of behaviour^38^. By merging both approaches, the rational inattention account predicts that low dopamine conditions will be associated with reductions in speed, speed-modulation *and* speed-meta-modulation.

Although our data can be considered consistent with the rational inattention account this, nevertheless, leaves unanswered questions about the exact mechanisms by which dopamine affects speed-modulation and speed-meta-modulation. The rational inattention account^37^ argues that dopamine ‘pays the cognitive costs’ of increasing the precision with which incoming information (e.g., trajectory information) is stored and represented but the exact nature of these ‘cognitive costs’ are yet to be determined. Mikhael and colleagues suggest attention as a candidate cost. If this were the case, in the context of our study, the hypothesis would be that in low dopamine conditions speed-modulation (and speed-meta-modulation) is reduced because participants attend less to changes in trajectory curvature because of a reluctance to pay the cognitive costs of selective attention. An alternative account has been forwarded by Manohar and colleagues^39^ who propose that, by signalling reward, dopamine permits more aggressive error correction. Thus, in our case, in low dopamine conditions, participants may attend to the changes in curvature but would nevertheless fail to appropriately modulate their speed due to a (presumably implicit) reluctance to pay the energetic costs of error correction. Further studies are required that specifically aim to tease apart whether, in low dopamine conditions, participants were less likely to attend to curvature changes, or whether they simply failed to adapt their movements to accommodate them.

Both Mikhael and colleagues and Manohar and colleagues suggest motivation-based mechanisms: They do not argue that dopamine changes one’s *ability* to pay attentional/effort-based costs, only one’s motivation to do so. Nevertheless, it is possible that dopamine plays a role in motor ability *per se*. Dopamine has long been linked to the invigoration of movements^40, 41^, is thought to play an important role in generating a range of different movement speeds, ^42^ and is key in signalling the start and end points of sub-movements. ^43^ Given the observation that distinct dopamine neurons are involved in reward signalling and self-paced movement^40^ (also see Engelhard et al. ^44^) it is feasible that dopaminergic manipulations directly affect participants’ ability to physically modulate their actions, independent of their motivation to do so. In other words, participants may be motivated to adapt movement speed as a function of curvature/global trajectory but may be unable to do so, perhaps because they have an inadequate range of movement speeds to choose from, and/or they are unable to signal the start/end points at which the adaptation should occur.

There are a number of possible reasons why the drug effect on speed-meta-modulation was observed only in Study 2 (haloperidol compared to placebo) and not in Study 1 (ON versus OFF dopaminergic medication in Parkinson’s). First, in Study 2 we were able to account for baseline striatal dopamine synthesis capacity by including working memory (a proxy measure of synthesis capacity) in our models, resulting in larger beta estimates for the effect of drug (0.107 compared to 0.008, see Supp Mats F). To be maximally sensitive to the effect of Parkinson’s medication, future studies investigating speed-meta-modulation should therefore endeavour to account for baseline striatal dopamine synthesis capacity. Secondly, given that Parkinson’s medication is thought to primarily boost tonic dopamine, ^35, 45^ whereas haloperidol affects both tonic and phasic dopamine, ^28, 46^ it is possible that speed-meta-modulation is influenced by phasic mechanisms that are much less restored by Parkinson’s medication. For example, recent models have posited a role for phasic dopamine in movement kinematics, specifically acting as part of a *“*velocity control circuit*”* ^47^. A final possibility is that different results in Study 1 and 2 are due to differences in the mechanisms of action of the two pharmacological interventions. Haloperidol primarily acts as a dopamine D2 receptor antagonist^46^ but can additionally affect cortical glutamate and noradrenaline function^48, 49^. Given that glutamate and noradrenaline have been implicated in prefrontal mechanisms underlying the flexible adaptation of on-going behaviour in response to environmental change^50-57^, and that speed-meta-modulation likely relies upon such mechanisms (e.g., changing behaviour to suit the current ‘environment’ (i.e., trajectory shape)), haloperidol’s effect on speed-meta-modulation may have been amplified by its modulation of other neurotransmitter systems in the prefrontal cortex. Whilst possible, this is perhaps unlikely given haloperidol’s high affinity with dopamine receptors relative to non-dopamine receptors.

Our research has practical implications for drug discovery and Parkinson’s treatment. Natural movements are made up of a wide range of trajectory shapes, and can be decomposed, using Fourier transform, into densities within different angular frequency bands. Thus, quantifying the effect of dopaminergic modulation across the angular frequency spectrum enables us to predict the effects of dopaminergic drugs on an extensive range of naturalistic movement trajectories. Future studies may build upon this to construct classification systems that can infer which drug an animal has taken based on their naturalistic movements^58^, and/or predict the effects of novel compounds (i.e., pharmacological interventions) on naturalistic movement.

To summarise, the current studies implicate dopamine in movement speed, speed-modulation and speed-meta-modulation for complex movements, thus extending the theoretical understanding of dopamine function beyond that which is conceptualised in current models of vigour.

## Methods

### Study 1

#### Participants

A total of 32 people with Parkinson’s were recruited, 19 male and 13 female (age range 46-78 years; mean (SD) = 63.2(7.6)). Mean number of years since diagnosis was 3.7(2.4) and the average Unified Parkinson’s Disease Rating Scale (Part II: Motor Aspects of Experiences of Daily Living; a self-report measure appropriate for assessing symptom severity^59^) was 11.91(6.93). All participants were taking a form of dopaminergic medication such as levodopa and/or dopamine agonists (see Supp Mats G), and their dosage was calculated as the levodopa equivalent dose using an online calculator (https://www.parkinsonsmeasurement.org/toolBox/levodopaEquivalentDose.htm).

Participants were recruited via Parkinson’s UK, the University of Birmingham Older Adults Database or social media. All participants gave fully informed consent and received renumeration of £10 per hour. The experimental procedure was approved by the local Research Ethics Committee (ERN_18-1800B).

#### Procedure

Participants completed two testing days, one to two days apart, which followed the same protocol but differed with respect to drug administration. On one of the days, participants completed the tasks prior to taking their first dose of dopaminergic medication in the morning. On the other day, participants completed the tasks approximately 1 hour after taking their first dose of dopaminergic medication. These protocols achieved ‘OFF’ and ‘ON’ medication states respectively and are standard practice in the literature. ^60-62^ The orders of the ON and OFF medication days were counterbalanced across participants. Following the medication protocol, participants undertook the shapes tracing task using a stylus and touch-screen device (Samsung Galaxy Tab A7). The task was programmed in PsychoPy and run on Pavlovia (PsychoJS platform version 2021.1.4).

##### Shapes tracing task

Participants traced four different shapes of varying angular frequencies (4/5, 4/3, 2, 4). The size of the shapes presented on the device did not exceed 9cm by 9cm. During each trial, participants were asked to trace around the shape on the screen ‘as fluidly as possible’ in a counter-clockwise direction, using a stylus held in their dominant hand. Participants completed eight blocks, two for each shape, presented in a random order. Each trial required participants to draw 10 full cycles of the shape, and the x and y coordinates of the stylus were recorded over time. During each block, participants had a total of seven attempts to complete four successful trials. Thus, a maximum of eight successful trials per shape was set, with participants limited to completing up to 14 trials to achieve this number. If participants significantly deviated from the shape (and into a region surrounding the shape which was not displayed to participants) or removed the stylus from the touch-screen (after a 5-second grace period), the trial was not classed as ‘successful’. Each trial had a timeout set at 90 seconds.

#### Data pre-processing

##### Shapes tracing task

Trials with fewer than 2.5 traces were excluded. The first samples of every trial which did not achieve a minimal speed (200 pixels per second), within the 5-second grace period, were removed to address a discontinuity of reported position at trial start. The maximum sampling rate of the tablet was 60Hz, but deviations occurred, thus positional data was resampled using the spline method to achieve a consistent 60Hz. The first ½ π angular displacement of each trial was discounted before data processing. Movement kinematics were calculated for each trial of the experiment using the x and y coordinates recorded over time. Movement speed was calculated as the first-order derivative of the positional non-null data. Maximum and minimum speed values were calculated as the average of the top and bottom 10% of speed values obtained during each trial. Speed-modulation values (also known as speed-curvature gradients; β in Equation 1) were calculated as the gradient of the relationship between movement speed (*v* in Equation 1) and the current curvature of the shape (*κ* in Equation 1), converted to an absolute value.

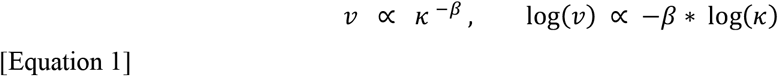

This followed the filtering and regression procedure set out in Huh & Sejnowski^11^ and Cook et al. ^18^, whereby log velocity was regressed against log curvature using MATLAB’s *regress* function to obtain speed-modulation values. Thus, a higher speed-modulation value indicates a steeper slope of the log speed to log curvature relationship. Speed-meta-modulation was calculated as the gradient of the regression line between speed-modulation values and an ordinal shape value (using MATLAB’s *polyfit* function to a polynomial power of 1). To meet normality assumptions, a log transform was applied to the movement speed data. Outliers of each dependent variable were removed, defined as values further than 2 standard deviations away from the mean.

##### Data subsetting

To ensure each participant had data across the range of shapes, speed-meta-modulation slopes were calculated from participants who had valid trials for at least the shapes with the highest and lowest angular frequency values (i.e., shapes 4/5 and 4 in Study 1). To keep analyses consistent, analyses on speed and speed-modulation in the main text were run on subsets of the data excluding participants who did not have valid trials for these shapes. This led to the removal of one ‘OFF medication’ dataset. Analyses on all datasets for speed and speed-modulation are reported in Supplementary Materials.

#### Analyses

All analyses were conducted in MATLAB 2022A. All mixed models were run using MATLAB’s *fitlme* function. Data and analysis scripts are available online at https://osf.io/vwu5t/?view_only=f1ce99b65142493bb313472f389c2e1f

##### Speed and Speed-modulation

To analyse the shapes tracing task data, effects-coded linear mixed models were employed for movement speed and speed-modulation values with drug state and shape as fixed effects, dosage interacting with drug state, and day, trial number and participant ID as random effects.

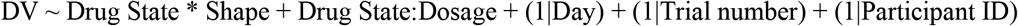

##### Maximum and Minimum Speed Values

Given the possible prediction highlighted in the Introduction that dopaminergic manipulation may disproportionately affect high speed movements, maximum and minimum speed values were analysed using the following LMM:

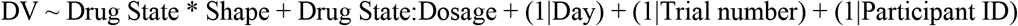

##### Speed-meta-modulation

For speed-meta-modulation, models were run using the following formula:

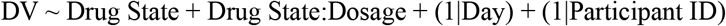

ANOVAs were conducted on the model coefficients to obtain p-values for the fixed effects.

### Study 2

#### Participants

43 participants (24 male, 19 female; age range 18-42 years, mean (SD) = 26.0(6.3)) passed a health screening prior to the experimental testing days following the protocol described in Rybicki et al. ^63^ and Schuster et al. ^29^ which involved Body Mass Index (BMI), blood pressure and electrocardiogram QT interval checks, as well as relevant medical history. Recruitment was conducted via the School of Psychology Research Participation Scheme or social media. All participants gave fully informed consent and received renumeration of £10 per hour. The experimental procedure was approved by the local Research Ethics Committee (ERN_18_1588).

#### Procedure

Participants completed two testing days, one to four weeks apart, lasting approximately 5.5 hours each. Following an on-the-day health check involving blood pressure and blood oxygenation levels, participants were administered capsules containing either 2.5mg of haloperidol (a dopamine D2 receptor antagonist) or a placebo, in a double-blind, placebo-controlled, within-subjects design. The orders of the days for haloperidol and placebo were pseudorandomised such that 50% of participants received haloperidol on day 1 and 50% on day 2. Haloperidol dosage and administration times were in line with previous studies in the literature demonstrating behavioural and psychological effects. ^28, 64^ At 1.75 hours post-tablet intake, participants completed a working memory task, followed by the shapes tracing task 4 hours post-tablet intake. Given that oral haloperidol is reported to be at its peak concentration in the blood plasma between 1.7 and 6.1 hours post-tablet intake on average^65^, both tasks were completed within the peak range of haloperidol blood plasma concentration, ensuring that drug action was likely to occur throughout administration of the tasks. Medical symptoms, blood pressure and mood were monitored before capsule administration, three times throughout the testing day, and at the end of the testing day.

##### Shapes Tracing Task

The general principles of the shapes tracing task remained the same as in Study 1, with a few exceptions. Three shapes were presented to participants (4/3, 2, and 4), as opposed to four shapes (Study 1: 4/5, 4/3, 2, and 4) to allow for a greater number of trials for the included shapes. The size of the shapes presented on the device did not exceed 12cm by 12cm. Each shape was traced for a total of 10 trials per shape, as opposed to a maximum of eight trials in Study 1, and each trial consisted of 10 x angular frequency (i.e., 10 × 2π radians of angular displacement) curvature oscillations (in the case of the ellipse and rounded square this is 10 complete traces of the shape). Participants were asked to repeat trials if they deviated from the shape or removed their stylus from the device. Participants could repeat the trial if they felt fluidity was not achieved (e.g., due to deviating from the shape or removing the stylus). The task was programmed and run using MATLAB 2014b 32-bit on a Surface Pro 4, using a touch-screen device to record participants’ movements (WACOM Cintiq 22 HD drawing tablet).

##### Working memory task

Participants completed a visual working memory task, adapted from the Sternberg visual working memory task^33^, programmed using MATLAB 2017b. The task involved 60 experimental trials across five blocks which were completed following 10 practice trials. In each trial, a fixation cross was presented in the centre of the screen (for a variable duration between 500-1000ms), followed by a list of letters (between 5 and 9 consonants depending on the block; 1000ms duration) and a blue fixation cross (3000ms duration). A single letter was then displayed (4000ms) and participants indicated with a keyboard press whether the letter was present in the previously displayed list (1 = yes, 2 = no, 3 = unsure). Accuracy and response time were recorded for each trial.

#### Data pre-processing

##### Shapes tracing task

As in Study 1 the first ½ π angular displacement of each trial was discounted before data processing. Movement speed (average, maximum and minimum values), speed-modulation values and speed-meta-modulation values were calculated for each trial using the x and y coordinates recorded over time, as in Study 1. To meet normality assumptions, a log transform was applied to the movement speed and speed-modulation data. Outliers of each dependent variable were removed, defined as values further than 2 standard deviations away from the mean. Three participants failed to complete both testing days, therefore two haloperidol datasets and one placebo dataset are missing from analyses.

##### Working memory task

As in previous studies^29, 63^, working memory span was calculated as the percentage of correct responses across all trials. Given evidence that baseline working memory span reliably predicts individual dopamine synthesis capacity^30, 31^, baseline striatal dopamine synthesis capacity was taken as the working memory span obtained under placebo. This value was used to explore any baseline dependent effects of the drug, as a body of literature reports that effects of dopaminergic drugs are modulated by striatal dopamine synthesis capacity. ^27-29^ Five participants failed to complete the working memory task at baseline, thus baseline striatal dopamine synthesis capacity could not be estimated and these participants could not be included in analyses incorporating this measure. We note that the correlation between working memory span and dopamine synthesis capacity was not replicated in a recent positron emission tomography (PET) imaging study^66^, but this may be due to the use of a less sensitive radioligand compared to earlier studies (18F-FDOPA rather than 18F-FMT).

##### Data subsetting

As in Study 1, speed-meta-modulation slopes were calculated from participants who had valid trials for at least the shapes with the highest and lowest angular frequency values (i.e., shapes 4/3 and 4 in Study 2), and analysis of the other two dependent variables was also conducted on this subset of participants. This reduced the sample size of analyses reported in the main text to 34 participants under placebo and 31 participants under haloperidol. Analyses on all datasets (N = 43) for speed and speed-modulation are reported in Supplementary Materials.

#### Analyses

All analyses were conducted in MATLAB 2022A. All mixed models were run using MATLAB’s *fitlme* function. As in Study 1, all linear mixed models were followed up with an ANOVA on the model coefficients to obtain p-values for the fixed effects. Data and analysis scripts are available online at https://osf.io/vwu5t/?view_only=f1ce99b65142493bb313472f389c2e1f

##### Speed and Speed-modulation

To analyse the dependent variables speed and speed-modulation, a series of effects-coded linear mixed models were employed. Reported in the main text is a mixed model that incorporated baseline striatal dopamine synthesis capacity as a fixed effect alongside drug state and shape, with day, trial number, and participant ID as random effects.

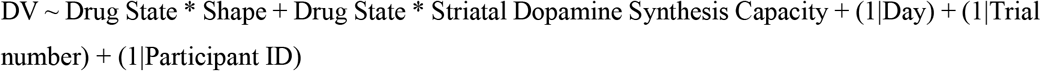

Supplementary Materials contain the results of mixed models without the incorporation of baseline striatal dopamine synthesis capacity (*Standard Models* in Supp Mats B (speed), Supp Mats D (speed-modulation) and Supp Mats F (speed-meta-modulation)).

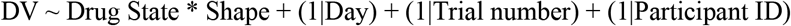

##### Maximum and Minimum Speed Values

Given the possible prediction highlighted in the discussion that dopaminergic manipulation may disproportionately affect high speed movements, maximum and minimum speed values were analysed using the following LMM:

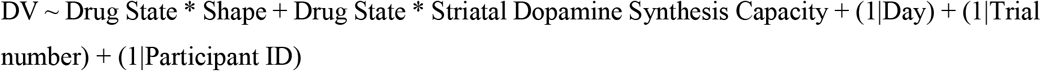

##### Speed-meta-modulation

For the speed-meta-modulation data, the following models were run, with (main text) and without (Supplementary Materials) striatal dopamine synthesis capacity respectively:

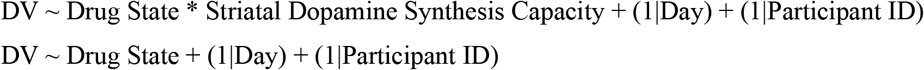

When a significant interaction between drug state and striatal dopamine synthesis capacity was found, this was unpacked by assessing the relationship between striatal dopamine synthesis capacity and the drug effect. For speed and speed-modulation, the drug effect was calculated for each participant as the mean value of the DV under placebo minus the mean value of the DV under haloperidol. For speed-meta-modulation, the drug effect was calculated as speed-meta-modulation under haloperidol minus speed-meta-modulation under placebo. Thus, for all DVs, positive values on the y-axis indicate reduced DV values under haloperidol. Following the drug effect calculation, a linear model was then employed with striatal dopamine synthesis capacity as predictor and Drug Effect and DV. To provide evidence in favour of an inverted-U function for the relationship between baseline dopamine and the dependent variable^32^, a negative linear relationship that cuts the x-axis would be expected when plotting the drug effect against estimated striatal dopamine synthesis capacity.

##### Independence of drug effects

Given the presence of drug effects on all three dependent variables in Study 2, further exploratory analyses were implemented on this dataset to investigate whether the drug effects are independent from each other. In each case, the control variable (e.g., speed) was included in a model predicting the dependent variable (e.g., speed-modulation), from which the residuals were saved. Models also included day as a random effect, and trial as a random effect for cases in which speed or speed-modulation were the dependent variable. A mixed model testing the effect of drug was run on the resultant residuals, using the corresponding model formula that included striatal dopamine synthesis capacity listed above.

## Acknowledgements

LJH was supported by a BBSRC PhD studentship provided by the BBSRC Midlands Integrative Biosciences Training Partnership [grant reference: BB/M01116X/1]. JLC was supported by the European Union’s Horizon 2020 Research and Innovation Programme under ERC-2017-StG Grant Agreement No. 757583 (Brain2Bee; Jennifer Cook PI). The project was further supported by a grant from the British Psychological Society Cognitive Section. We thank our lived experience experts for reviewing the project, including Mary Adcock, Penny King, Rebecca Millward, and Sara Moore, and Parkinson’s UK for their assistance with participant involvement and recruitment.

## Data and Code Availability

Data and analysis scripts are available online at https://osf.io/vwu5t/?view_only=f1ce99b65142493bb313472f389c2e1f

## Notes

**Disclosure of Interest** The authors report no conflict of interest.

https://osf.io/vwu5t/?view_only=f1ce99b65142493bb313472f389c2e1f

### Competing Interest Statement

The authors have declared no competing interest.

