## Supplementary Materials for "Dopaminergic manipulations affect the modulation and meta-modulation of movement speed: evidence from two pharmacological interventions"

#### Supplementary Materials A

***Removal of Parkinson's medication reduces movement speed.***

*Additional Mixed Model Details*

To explore the main effect of shape on movement speed, post-hoc tests were conducted on subsets of the data to compare two levels of shape. These tests revealed that the overall main effect of shape reflected lower movement speeds for lower angular frequency shapes. Shape 4/5 was traced with the slowest movement speed, followed by shape 4/3 (main effect of shape between shapes 4/5 and 4/3: $F(1, 963) = 127.95$, $p < .001$), followed by shape 4 (main effect of shape between shapes 4/3 and 4: $F(1, 941) = 18.50$, $p < .001$), with shape 2 traced with the highest speed (main effect of shape between shapes 4 and 2: $F(1, 921) = 28.91$, $p < .001$). The interaction between drug state and shape was non-significant ($F(3, 1885) = 0.29$, $p = .832$) as was the interaction between drug state and dosage ($F(1, 1885) = 0.15$, $p = .699$).

*Analysis on All Data*

Analyses reported in the main text (which were conducted on a dataset in which all participants had valid trials for at least the shapes with the highest and lowest angular frequency values) were repeated for a dataset containing all data, following the removal of outliers and a log transform. Importantly our core finding - a main effect of drug state - was observed ($F(1, 1905) = 71.79$, $p < .001$), with lower movement speeds observed OFF medication (beta estimate = -0.078, 95% CI [-0.096, -0.060]). As in our primary analysis reported in the main text, a main effect of shape was observed ($F(3, 1905) = 137.96$, $p < .001$), with no drug by shape interaction ($F(3, 1905) = 0.33$, $p = .804$) or drug by dosage interaction ($F(1, 1905) = 0.09$, $p = .762$).

#### Supplementary Materials B

***Haloperidol reduces movement speed.***

*Additional Mixed Model Details*

The main effect of shape on movement speed reflected lower movement speeds for lower angular frequency shapes. As revealed by post-hoc tests assessing the main effect of shape between two levels of the condition, shape 4/3 was traced with the slowest movement speed, followed by shape 4 (main effect of shape between shapes 4/3 and 4: $F(1, 1111) = 15.03$, $p < .001$), with shape 2 traced with the highest speed (main effect of shape between shapes 4 and 2: $F(1, 1039) = 217.10$, $p < .001$). The interaction between drug state and shape was non-significant ($F(2, 1600) = 2.79$, $p = .062$).

*Standard Model (not including estimated baseline striatal synthesis capacity)*

A LMM including drug state and shape as fixed effects (but not estimated baseline striatal synthesis capacity) revealed a significant main effect of drug state ($F(1, 1700) = 10.34$, $p = .001$), with lower movement speed values in the haloperidol condition compared to the placebo condition (beta estimate: -0.023, 95% CI [-

0.037, -0.009]). As in our primary analysis, we observed a significant main effect of shape ($F(2, 1700) = 157.20$, $p < .001$). Unpacking this further with post-hoc tests revealed that, as in the model above, shape 4/3 was traced with the slowest movement speed, followed by shape 4 (main effect of shape between shapes 4/3 and 4: $F(1, 1177) = 9.96$, $p = .002$), followed by shape 2 (main effect of shape between shapes 4 and 2: $F(1, 1105) = 216.97$, $p < .001$). In addition, we observed an interaction between drug state and shape ($F(2,1700) = 4.20$, $p = .015$) in which the strongest drug effect was observed for shape 4/3 ($F(1, 595) = 6.53$, $p = .011$, beta estimate = -0.024, 95% CI [-0.042, -0.005]), followed by shape 2 ($F(1, 523) = 6.04$, $p = .014$, beta estimate = -0.033, 95% CI [-0.060, -0.007]), followed by shape 4 ($F(1, 582) = 1.57$, $p = .211$, beta estimate = -0.012, 95% CI [-0.030, 0.007]). As such, we again present no evidence that shapes typically traced at higher speeds are disproportionately affected by the drug (as shape 4/3 was traced the slowest but had the largest drug effect).

*Analysis on All Data*

Analysis reported in main text was repeated for a dataset containing all data, following the removal of outliers and a log transform. An LMM including drug state, shape and estimated baseline striatal synthesis capacity as fixed effects revealed a significant main effect of drug state ($F(1, 1673) = 46.73$, $p < .001$) with lower movement speed values in the haloperidol condition compared to the placebo condition (beta estimate: -0.505, 95% CI [-0.650, -0.360]). Again, a main effect of shape ($F(2, 1673) = 153.39$, $p < .001$) was identified, which reflected a pattern whereby shape 4/3 was traced the slowest, followed by shape 4 (main effect of shape between shapes 4/3 and 4: $F(1, 1163) = 12.89$, $p <.001$), followed by shape 2 (main effect of shape between shapes 4 and 2: $F(1, 1085) = 176.04$, $p < .001$). We also observed an interaction between drug state and shape ($F(2, 1673) = 3.30$, $p = .037$), whereby the largest drug effect was observed for shape 4/3 ($F(1, 586) = 52.56$, $p < .001$, beta estimate = -0.712, 95% CI [-0.905, -0.519]), followed by shape 4 ($F(1, 575) = 20.05$, $p < .001$, beta estimate = -0.438, 95% CI [-0.629, -0.246]), followed by shape 2 ($F(1, 508) = 6.73$, $p = .010$, beta estimate = -0.392, 95% CI [-0.688, -0.095]). Again, this pattern did not provide evidence that shapes traced at higher speeds are disproportionately affected by the drug. The overall main effect of drug state was moderated by estimated striatal dopamine synthesis capacity ($F(1,1673) = 43.21$, $p <.001$), and again a negative linear relationship was found between drug effect and estimated striatal dopamine synthesis capacity which cut the x-axis ($F(1, 29) = 4.68$, $p = .039$; beta estimate = -0.016, 95% CI [-0.032, -0.001]). Thus, with the full dataset the pattern of results is the same as in our primary analysis but with the addition of a significant interaction between drug state and shape.

An LMM including drug state and shape as fixed effects (but not estimated baseline striatal synthesis capacity) revealed a significant main effect of drug state ($F(1, 1773) = 11.05$, $p = .001$), with lower movement speed values in the haloperidol condition compared to the placebo condition (beta estimate: -0.022, 95% CI [-0.035, -0.009]). We observed a significant main effect of shape ($F(2, 1773) = 137.50$, $p < .001$), with shape 4/3 traced the slowest, followed by shape 4 (main effect of shape between shapes 4/3 and 4: $F(1, 1229) = 8.18$, $p = .004$), followed by shape 2 (main effect of shape between shapes 4 and 2: $F(1, 1151) = 176.97$, $p$

< .001). The interaction between drug state and shape was also significant ($F$(2,1773) = 3.88, $p$ = .021), with the strongest drug effect of shape 4/3 ($F$(1, 622) = 7.70, $p$ = .006, beta estimate = -0.024, 95% CI [-0.041, -0.007]), followed by shape 2 ($F$(1, 544) = 3.52, $p$ = .061, beta estimate = -0.025, 95% CI [-0.052, 0.001]), followed by shape 4 ($F$(1, 607) = 0.47, $p$ = .493, beta estimate = -0.006, 95% CI [-0.023, 0.011]). This pattern of results, again, does not indicate that shapes traced at faster speeds are disproportionally affected by the drug.

**Supplementary Materials C**

***Removal of PD medication affects speed-modulation.***

*Additional Mixed Model Details*

Post-hoc tests were conducted on subsets of the data to explore the main effect of shape on speed modulation, whereby the main effect of shape for pairs of the shape condition were assessed. These analyses revealed that the main effect of shape on speed modulation (F(3, 1906) = 676.08, p < .001) reflected a pattern in which shape 4 was traced with the lowest speed modulation values and shape 2 was traced with the highest, with shape 4/3 falling below shape 2 (main effect of shape between shapes 4/3 and 2: $F$(1, 951) = 115.36, $p$ < .001), followed by shape 4/5 (main effect of shape between shapes 4/5 and 4/3: $F$(1, 933) = 39.31, $p$ < .001; main effect of shape between shapes 4/5 and 4: $F$(1, 954) = 695.78, $p$ < .001). The interaction between drug state and dosage was non-significant ($F$(1, 1906) = 2.89, $p$ = .089).

*Analysis on All Data*

Analysis reported in the main text was repeated for a dataset containing all data, following the removal of outliers and a log transform. As in our primary analysis, a main effect of drug state was observed ($F$(1, 1925) = 7.18, $p$ = .007), with lower speed modulation values observed OFF medication (beta estimate = -0.016, 95% CI [-0.028, -0.004]). Again, a main effect of shape was observed ($F$(3, 1925) = 672.14, $p$ < .001), and no drug by dosage interaction ($F$(1, 1925) = 2.67, $p$ = .102).

**Supplementary Materials D**

***Haloperidol affects speed-modulation.***

*Additional Mixed Model Details*

As indicated by post-hoc analyses on subsets of the data, the main effect of shape on speed modulation ($F$(2, 1642) = 1092.40, $p$ < .001) reflected a pattern in which shape 4 was traced with the lowest speed modulation values and shape 4/3 was traced with the highest, with shape 2 falling between shapes 4/3 and 4 (main effect of shape between shapes 4/3 and 2: $F$(1, 1106) = 8.32, $p$ = .004; main effect of shape between shapes 2 and 4: $F$(1, 1085) = 1540.70, $p$ < .001).

*Standard Model (not including estimated baseline striatal synthesis capacity)*

A LMM including drug state and shape as fixed effects revealed a significant main effect of drug state ($F(1, 1730) = 29.85$, $p < .001$; beta estimate = 0.020, 95% CI [0.013, 0.027]). Again, a main effect of shape was observed ($F(2, 1730) = 1036.60$, $p < .001$).

*Analysis on All Data*

Analysis reported in the main text was repeated for a dataset containing all data, following the removal of outliers and a log transform. An LMM including drug state, shape and estimated baseline striatal synthesis capacity as fixed effects revealed a significant main effect of drug state ($F(1, 1719) = 48.10$, $p < .001$) with lower speed modulation values in the haloperidol condition compared to the placebo condition (beta estimate: -0.256, 95% CI [-0.329, -0.184]). Again, a main effect of shape ($F(2, 1719) = 1127.40$, $p < .001$) was identified. The main effect of drug state was moderated by estimated striatal dopamine synthesis capacity ($F(1,1719) = 54.82$, $p <.001$), and again a negative linear relationship was found between drug effect and estimated striatal dopamine synthesis capacity which cut the x-axis ($F(1, 29) = 7.41$, $p = .011$; beta estimate = -0.007, 95% CI [-0.013, -0.002]). Thus, this analysis on the full dataset revealed the same pattern of results as in our primary analysis.

An LMM including drug state and shape as fixed effects revealed a significant main effect of drug state ($F(1, 1807) = 18.81$, $p < .001$; beta estimate = 0.015, 95% CI [0.008, 0.022]). Again, a main effect of shape was observed ($F(2, 1807) = 1065.80$, $p < .001$).

**Supplementary Materials E**

***Removal of PD medication does not affect speed-meta-modulation.***

*Additional Mixed Model Details*

No main effect of drug state was observed ($F(1, 57) = 0.49$, $p = .485$), nor an interaction between drug state and dosage ($F(1, 57) = 1.04$, $p = .313$). This lack of drug effect on speed meta-modulation is supported by the lack of a drug state by shape interaction in the speed-modulation LMM for Study 1 ($F(3, 1906) = 0.30$, $p = .824$).

**Supplementary Materials F**

***Haloperidol affects speed-meta-modulation.***

*Standard Model (not including estimated baseline striatal synthesis capacity)*

When analysing the data without including estimated baseline striatal synthesis capacity as a predictor, the main effect of drug state on speed-meta-modulation remained. We again observed that haloperidol reduced the meta-modulation of speed ($F(1,59) = 7.93$, $p = .007$; beta estimate = 0.008 95% CI [0.002, 0.013]), with shallower meta-modulation gradients under haloperidol.

### Supplementary Materials G

***Table S1 Types of medications taken by participants in Study 1.***

| | **N** |
|---|---|
| **Combined Totals** | |
| Levodopa | 23 |
| Dopamine Agonists | 16 |
| MAO Inhibitors | 10 |
| **Those Taking One Type of Medication** | |
| Levodopa | 9 |
| Dopamine Agonists | 5 |
| MAO Inhibitors | 2 |
| **Those Taking Two Types of Medications** | |
| Levodopa & Dopamine Agonists | 11 |
| Levodopa & MAO Inhibitors | 2 |
| Dopamine Agonists & MAO Inhibitors | 1 |
| **Those Taking Three Types of Medications** | |
| Levodopa, Dopamine Agonists & MAO Inhibitors | 2 |
